# Neural Representation of Associative Threat Learning in Pulvinar Divisions, Lateral Geniculate Nucleus, and Mediodorsal Thalamus in Humans

**DOI:** 10.1101/2025.07.09.663823

**Authors:** Muhammad Badarnee, Zhenfu Wen, B. Isabel Moallem, Stephen Maren, Mohammed R Milad

## Abstract

Understanding the neural mechanisms underlying associative threat learning is essential for advancing behavioral models of threat and adaptation. We investigated distinct activation patterns across thalamic pulvinar divisions, the lateral geniculate nucleus (LGN), and the mediodorsal thalamus (MD) during associative threat learning using fMRI. The anterior pulvinar and MD exhibited parallel activation patterns, that may reflect distinct contributions to automatic and more deliberative learning processes. Additionally, our findings suggest a hierarchical functional organization of pulvinar activation during fear conditioning, in which coordinated activation among inferior, lateral, medial, and anterior divisions may support the integration of threat-related information. Pulvinar divisions and the MD showed activation during extinction learning and exhibited activation patterns consistent with salience processing during extinction recall and threat renewal. LGN activation patterns were consistent with feedforward visual processing. These findings extend current models of threat learning and memory by suggesting distinct patterns of thalamic involvement across associative threat learning and memory and providing a functional framework for investigating thalamic contributions to these processes.

## Introduction

Over a century ago, Ivan Pavlov provided foundational behavioral evidence for associative learning, demonstrating that pairing a neutral stimulus with an unconditioned stimulus could elicit a conditioned response (Pavlov, 1904). This discovery laid the groundwork for modern learning theories and has been particularly influential in understanding how humans learn to associate neutral stimuli with threats, a process known as associative threat learning. This survival-oriented learning plays a key role in shaping a broad range of behavioral and emotional responses, such as avoidance, decision-making under threats, and fear regulation (Badarnee et al., 2025; Kolling et al., 2014; Korn and Bach, 2019, 2018; Milad et al., 2013). Research on the neural mechanisms underlying associative threat learning has primarily focused on brain regions involved in emotional regulation, such as the amygdala, hippocampus, and prefrontal cortex (PFC) (Fullana et al., 2015; Krasne et al., 2021; Milad and Quirk, 2012; Vuilleumier et al., 2003). More recently, the thalamus has received growing attention in threat learning (Lithari et al., 2015; Penzo et al., 2015; Ramanathan et al., 2018; Ramanathan and Maren, 2019; Ratigan et al., 2023; Totty et al., 2023), since its role has been reconsidered beyond the traditional view of a mere relay station. Yet, the distinct contribution of this complex structure to associative threat learning remains poorly understood.

The thalamus is a hub structure in the mammalian brain that plays multiple critical roles, ranging from basic sensory processing to higher-order functions and threat learning (Halassa and Sherman, 2019; Hummos et al., 2022; Hwang et al., 2017; Saalmann and Kastner, 2011; Sherman, 2016, 2007). It has long been proposed that threat detection is mediated by the pulvinar and the lateral geniculate nucleus (LGN) (Carr, 2015; Pessoa and Adolphs, 2010; Silverstein and Ingvar, 2015), two major nuclei of the visual thalamus (Arcaro et al., 2015; Casanova and Chalupa, 2023; Takakuwa et al., 2021). The pulvinar projects directly to the amygdala and is believed to subconsciously facilitate rapid survival reactions to threats via a subcortical path (‘low road’) (Carr, 2015; Pessoa and Adolphs, 2010; Rafal et al., 2015; Wei et al., 2015). The LGN, on the other hand, is part of a slower but more accurate neural pathway, connecting peripheral information to cortical regions for comprehensive processing (‘high road’). Additionally, the involvement of the mediodorsal thalamus (MD) in threat learning has also been proposed(Lee and Shin, 2016; Lee et al., 2011). Although this nucleus is not typically classified as part of the visual thalamus, it serves as a high-order region reciprocally connected to the amygdala and PFC while playing a critical role in executive functions (Hwang et al., 2020; Li et al., 2022; Mukherjee et al., 2021; Wolff and Halassa, 2024).

The general implication of these nuclei in threat processing has been demonstrated in primates and rodent models. It has been shown that exposing primates to snake images elicited increased neuronal firing within the pulvinar (Le et al., 2013). Excitation of neurons within the LGN enhanced the acquisition of eyeblink conditioning in rodents (Halverson and Freeman, 2010). Inhibiting this nucleus was associated with impaired conditioned responses (Shi and Davis, 2001; Steinmetz et al., 2013), and activating GABA neurons in the LGN reduced freezing response to an overhead dark shadow that mimics a real-world predator in rats (Salay and Huberman, 2021). Reduced freezing was also observed after MD lesions (Li et al., 2004) and when the connections with the anterior cingulate cortex were ablated (Zheng et al., 2020). Lesions within the MD are also associated with impaired fear extinction (Lee et al., 2011), and injecting gabazine, a modulator of extra-synaptic GABA receptors, into the MD facilitated extinction (Paydar et al., 2014).

Evidence regarding the involvement of the thalamus in the context of threat learning in humans is relatively sparse. Much of our knowledge comes from studies focusing on attention, perception, and decision-making. For example, the LGN has been mainly viewed as a transmitter of peripheral information to other brain regions and found to be associated with selective attention and anticipation of visual stimuli (Mahoney and Schmidt, 2024; O’Connor et al., 2002; Saalmann and Kastner, 2009). The MD contribution to perception, spatial attention, and decision-making has also been reported (Griffiths et al., 2022; Wurtz et al., 2011). Beyond cognitive research, some human studies have specifically investigated the pulvinar’s engagement in threat detection. Individuals with increased fiber density in the pulvinar-amygdala pathway showed an enhanced ability to recognize fearful faces (McFadyen et al., 2019). A patient with a complete lesion in the left pulvinar showed slower responses to threatening images when stimuli were presented on the ipsilesional, but not contralesional, field (Ward et al., 2005). In agreement with this, unseen stimuli (fearful vs. happy faces) presented on the blind side of patients with hemianopia moderated their performance when the pulvinar was spared but not when it was lesioned (Bertini et al., 2018).

Specific thalamic contribution to threat processing is still largely unknown, and translational research on this topic is limited. We aim to address these critical gaps in our understanding by investigating the distinct neural representations of associative threat learning in the human pulvinar divisions, LGN, and MD. We analyzed the neuroimaging data of 293-412 controls. All participants underwent a two-day threat learning paradigm (Milad et al., 2009, 2007; Wen et al., 2024, 2022) while in the fMRI scanner. The conditioned stimulus (CS+) (e.g., red light) was associated with an electric shock (unconditioned stimulus US, 62.5% reinforcement rate), while a control light (e.g., blue) was never paired with the US (CS-). This conditioning phase occurred in a computer-displayed visual context A (e.g., an office). The extinction learning included presenting the CS+ and CS- with no US reinforcement in a distinct context B (e.g., bookcase). In extinction recall and threat renewal, the extinguished stimuli were presented with no US reinforcement within contexts B (safe contextual cues) and A (threat contextual cues), respectively. Schematic illustrations of each phase of the paradigm are presented in **Figs. 1a, 4a-6a**.

**Fig. 1.**
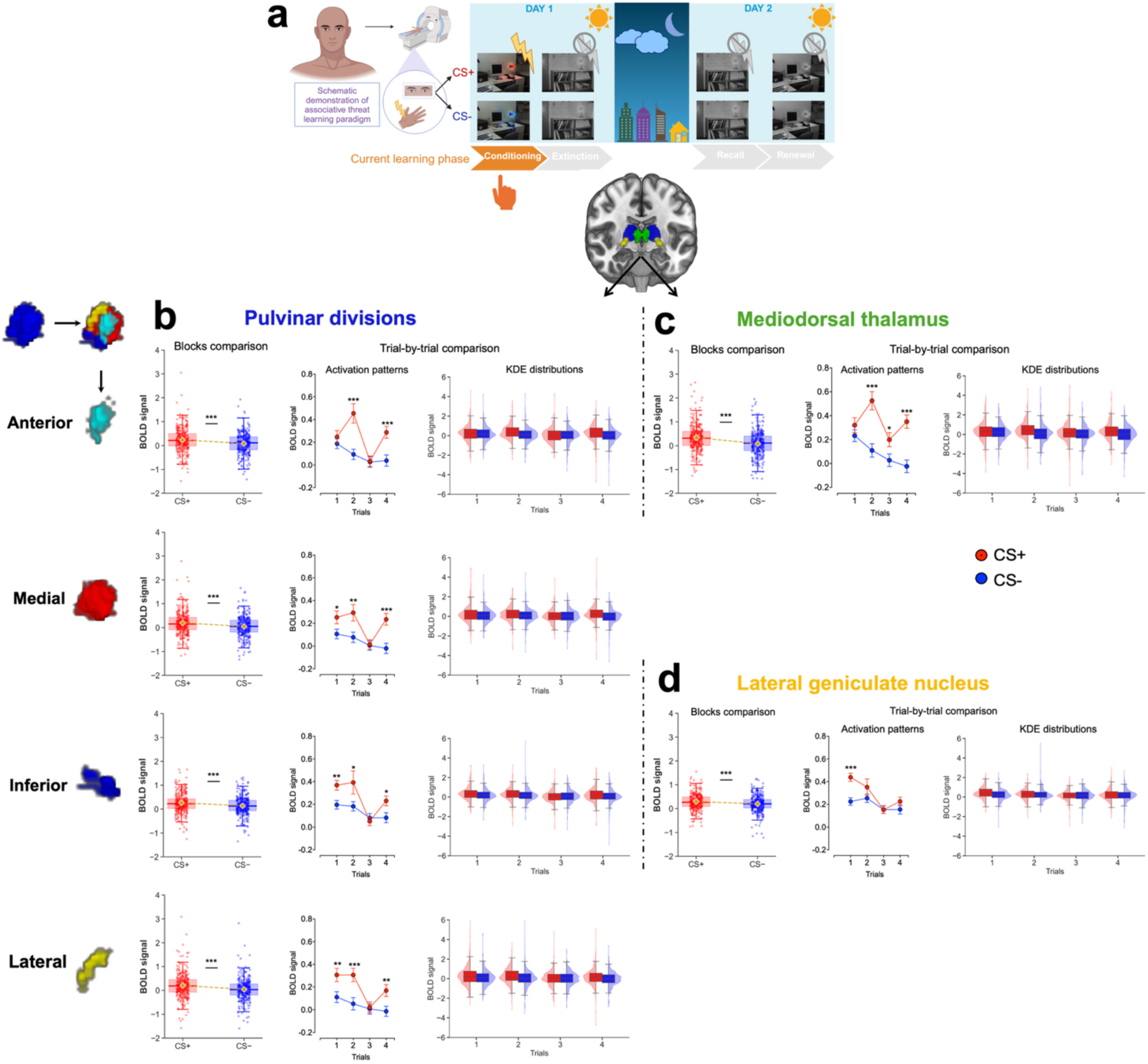
Neural representation during associative threat learning in pulvinar divisions, MD, and LGN. (a) Human fear conditioning paradigm during fMRI. (b-d) Boxplots with individual data points at the block level (gold diamonds indicate the means), and trial-wise activation shown as means ± SE and kernel density estimates (KDEs) for responses to CS+ and CS- in the pulvinar divisions (b), MD (c), and LGN (d). Block-level comparisons were assessed using paired *t*-tests, while trial-level effects were examined using a 2 × 2 repeated-measures ANOVA, followed by post hoc comparisons between CS+ and CS− across the four trials. Multiple comparisons were controlled using false discovery rate (FDR) correction. Conditioning sample size: *n* = 293. Detailed statistical parameters are provided in **Supplementary Tables 1-2**. Additionally, cross-validation analyses were conducted to assess the robustness of the CS+ vs. CS− signal in the anterior pulvinar and MD (**Supplementary Figs. 1-2** and **Supplementary File 1**). *p < 0.05, **p < 0.01, ***p < 0.001. SE, standard error; KDE, kernel density estimates; MD, mediodorsal thalamus; LGN, lateral geniculate nucleus; CS+, conditioned stimulus predicting shock; CS−, conditioned stimulus predicting no shock. Display items in panel (a) were created using BioRender (BioRender.com/t54zvyf).

We focus on neural activation patterns within pulvinar divisions, LGN, and MD while acquiring the CS-US association. As associative learning is a rapid psychological process (Konrad et al., 2024), we analyzed the first four trials of all threat learning phases. The purpose of this study is to investigate thalamic function during each learning phase separately, focusing on CS+ vs. CS− differences within phases rather than comparing activation across phases. This phase-specific approach allows us to characterize thalamic functional dynamics within each stage of learning and memory, avoiding potential confounds arising from the distinct processes of conditioning, extinction, and recall. We compared brain activation (BOLD) to the CS+ vs. CS- at the block level by averaging the activation across all four trials and at a trial-by-trial level to provide finer activation temporal resolution. This dual approach allows us to capture the general neural responses associated with associative threat learning and provides new insights into trial-level dynamics (Wen et al., 2022). The current dominant neurocognitive thalamic models (Sherman, 2007; Sherman and Guillery, 2006) highlight the thalamus’s role in mediating cortical-cortical communications and facilitating high-order functions. Based on this framework, we anticipated distinct activation patterns across the pulvinar, MD, and LGN during associative threat learning. Given its established role as a first-order thalamic nucleus, the LGN was expected to show activation patterns distinct from those of the higher-order pulvinar and MD.

## Results

### Parallel activation of the anterior pulvinar and MD during conditioning is consistent with associative threat learning

At the block level, both the anterior pulvinar and MD showed increased activation to CS+ vs. CS− (anterior pulvinar: *t_(292)_ = 4.41, p = 0.00001, d = 0.25*; MD: *t_(292)_ = 6.41, p = 5.83x10^-10^, d = 0.37*; **Fig. 1b–c**), suggesting a possible involvement of these regions in early associative threat learning. The trial-wise analysis revealed a distinct activation pattern. On the first trial, activation did not differ between CS+ and CS− in either the anterior pulvinar (*p_FDR_ = 0.44*) or the MD (*p_FDR_ = 0.20*). In contrast, on the subsequent trial, both regions showed significantly greater BOLD responses to CS+ than CS− (anterior pulvinar: *p_FDR_ = 0.000001*; MD: *p_FDR_ = 0.000003*; **Fig. 1b–c**; Detailed statistical parameters are provided in **Supplementary Tables 1–2**). In our paradigm, the electric shock was paired with the CS+ at the end of the CS presentation. The similar BOLD responses to the CS+ and CS- during the first trial may reflect an initially similar neural response to the two stimuli before the CS-US association was acquired. The subsequent increase in activation for the CS+ by the second trial is consistent with rapid associative learning and may reflect changes in the neural representation of the conditioned stimulus.

The similarity in trial-level activation patterns in the anterior pulvinar and MD raises the question of whether this apparent similarity consistent with a parallel functional contribution to associative threat learning. To test this possibility, we first quantified the trial-wise relationships using Pearson correlation coefficients between the two regions. The observed correlations support consistent co-activation across all corresponding trials (*r values ≥ 0.66, p < 0.001*; **Fig. 2a**, left). However, this approach could not exclude the possibility that the shared variance is a consequence of shared anatomical proximity, particularly because both regions are part of the same brain structure, i.e., the thalamus. To control for this confound, we conducted a hierarchical regression model in four steps for each trial. We modeled MD activation as the target and progressively added the pulvinar divisions as covariates. Despite minor variations in regression coefficients across control models, the anterior pulvinar consistently showed the largest standardized regression coefficients (all *p < 0.01;* **Fig. 2a**, middle) and accounted for a substantial proportion of the explained variance in MD activation (*R² = 0.43-0.61*). In contrast, other pulvinar divisions contributed minimally to the explained variance (*across all other pulvinar divisions, the maximum observed ΔR² was 0.02* in trial 1, *0.01* in trial 2, *0.01* in trial 3, and *0.002* in trial 4; **Fig. 2a**). These results were stable across 10,000 bootstrap resamples with replacement, indicating that the association between anterior pulvinar activation and MD activation is robust to resampling variability (**Fig. 2a**, right; additionally, see alternative hypothesis tests of the anterior pulvinar-MD relationships in Supplementary Fig. 3).

**Fig 2.**
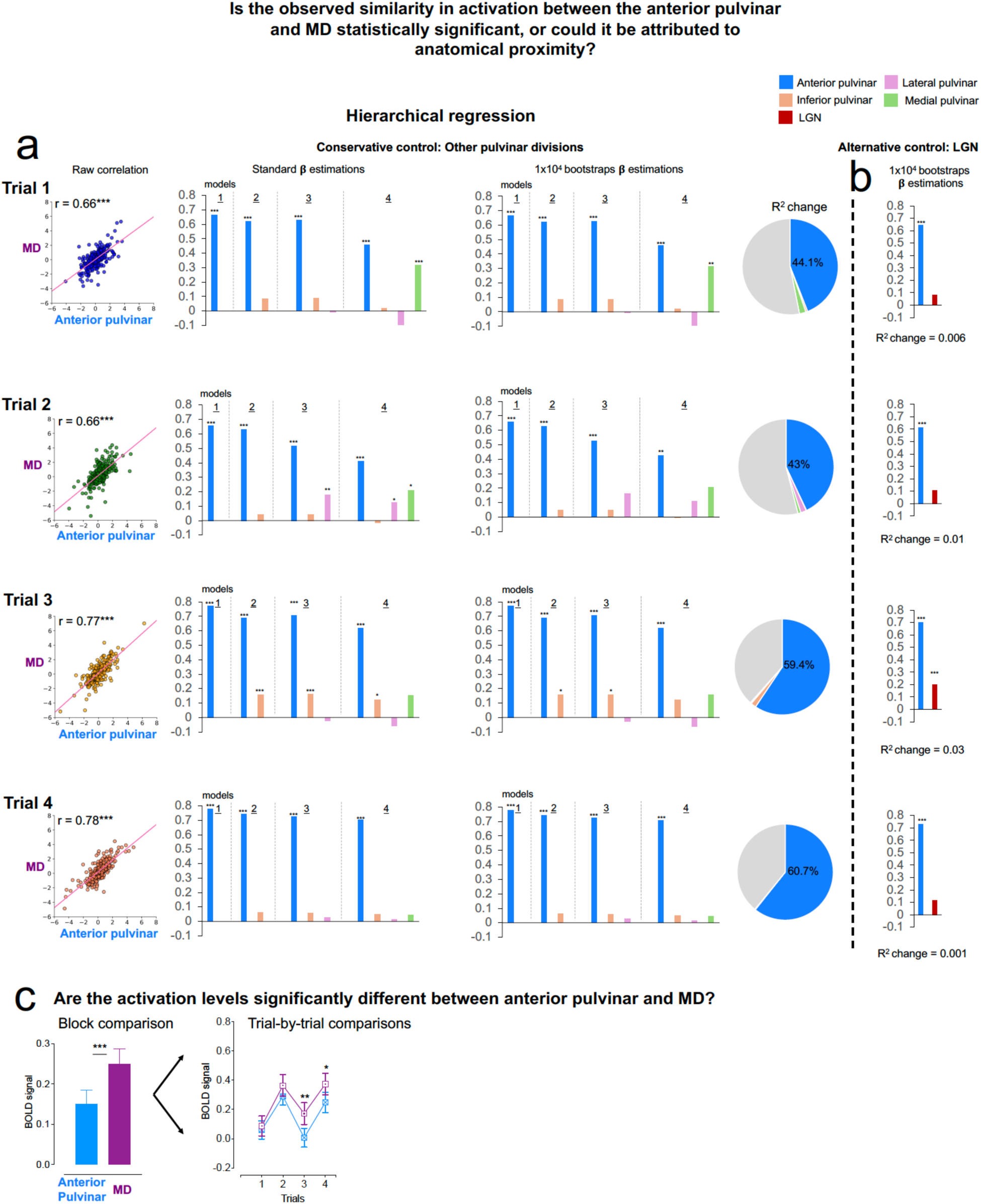
Quantifying the relationships between the anterior pulvinar and MD during conditioning. **a.** Hierarchical regression models of trial-wise relationships between the anterior pulvinar and MD activations while controlling for a potential effect of anatomical proximity. The effect of anatomical proximity was controlled by progressively adding other pulvinar divisions as controls. **b.** Results of a hierarchical model that included the LGN as an alternative control, distinct from the pulvinar. **c.** Comparison of activation levels in the anterior pulvinar and MD at both block-wise and trial-wise levels (means ± SE). Sample size in these analyses: *n* = 293. Supplementary analyses testing alternative hypotheses regarding the correlations between the MD and all pulvinar divisions are presented in **Supplementary Fig. 3**. *p<0.05, **p<0.01, ***p<0.001 MD: Mediodorsal thalamus. LGN: Lateral geniculate nucleus. SE: Standard error.

Applying the same analytical approach while additionally controlling for LGN activation as an anatomical control, beyond the pulvinar itself, yielded comparable results. Anterior pulvinar activation continued to account for the majority of the explained variance in MD activation, whereas the inclusion of LGN activation produced only an incremental additional increase in explained variance (*ΔR² = 0.006* in trial 1, *0.01* in trial 2, *0.03* in trial 3, and *0.01* in trial 4; **Fig. 2b**). Together, these analyses support preferential trial-wise functional co-activations between the anterior pulvinar and MD, independent of shared anatomical proximity.

Although co-activation does not necessarily imply similar activation magnitude, we further tested the differences in activation levels in the anterior pulvinar and MD. We used a t-test to capture the differences in the overall activation at the block level and repeated measures analysis of variance (RM-ANOVA) to capture the activation differences at the trial level. The results showed that the overall MD activation in response to CS+ was higher than the anterior pulvinar response (*t_(292)_ = 3.49, p = 0.0005, d = 0.20*, **Fig. 2c**). The trial-wise analysis revealed that the overall differences between the two regions is driven particularly from the activation in trials 3 and 4 (*F_(1, 292)_ = 5.84, p = 0.01; Trial 1: t_(292)_ = 0.49, p_FDR_ = 0.82, d = 0.02; Trial 2: t_(292)_ = 0.03, p_FDR_ = 0.97, d = 0.002, Trial 3: t_(292)_ = 3.40, p_FDR_ = 0.003, d = 0.199; Trial 4: t_(292)_ = 2.63, p_FDR_ = 0.01, d = 0.15;* **Fig. 2c**). Although the anterior pulvinar and MD exhibited similar trial-wise activation patterns, the greater activation observed in the MD suggests that these regions may differ in the extent of their engagement during associative threat learning.

### A data-driven approach suggests a hierarchical computational organization of pulvinar divisions during associative threat learning

The anatomical and functional interconnections between pulvinar divisions remain insufficiently characterized. We integrated findings from complementary analyses to propose a computational framework describing the functional relationships among pulvinar divisions during associative threat learning.

We first defined the activation within each pulvinar division as the differences in BOLD signal at the block level in response to CS+ compared to CS-. We then performed RM-ANOVA to test whether the pulvinar divisions were engaged with different activation levels. We found no activation differences, suggesting similar processing levels of CS+ information across all pulvinar divisions (*F_(2.29, 670.33)_ = 0.96, p = 0.39*; **Fig. 3a**). Using network analysis, we explored the underlying functional dynamics between the divisions. We employed the EBICglasso method to estimate a sparse Gaussian graphical model (Foygel and Drton, 2010). To assess the stability of the network and provide robust estimates of edges and centrality measures, we conducted 10,000 bootstrap resamples. This approach allowed us to compute 95% confidence intervals (CI), ensuring the reliability of our findings. The combined use of EBICglasso and bootstrapping is essential for accurately capturing the dynamics of pulvinar division interactions. The resulting network included four nodes and five edges. The network plot identified a statistical association between the lateral and anterior pulvinar, together with pathways linking the lateral and inferior pulvinar to the anterior pulvinar through the medial pulvinar (**Fig. 3b**). Additionally, the medial pulvinar exhibited the highest centrality values, a measure of node importance within the network, consistent with the possibility that this division services a hub-like position within the identified functional network during associative threat learning (**Fig. 3b**).

**Fig. 3.**
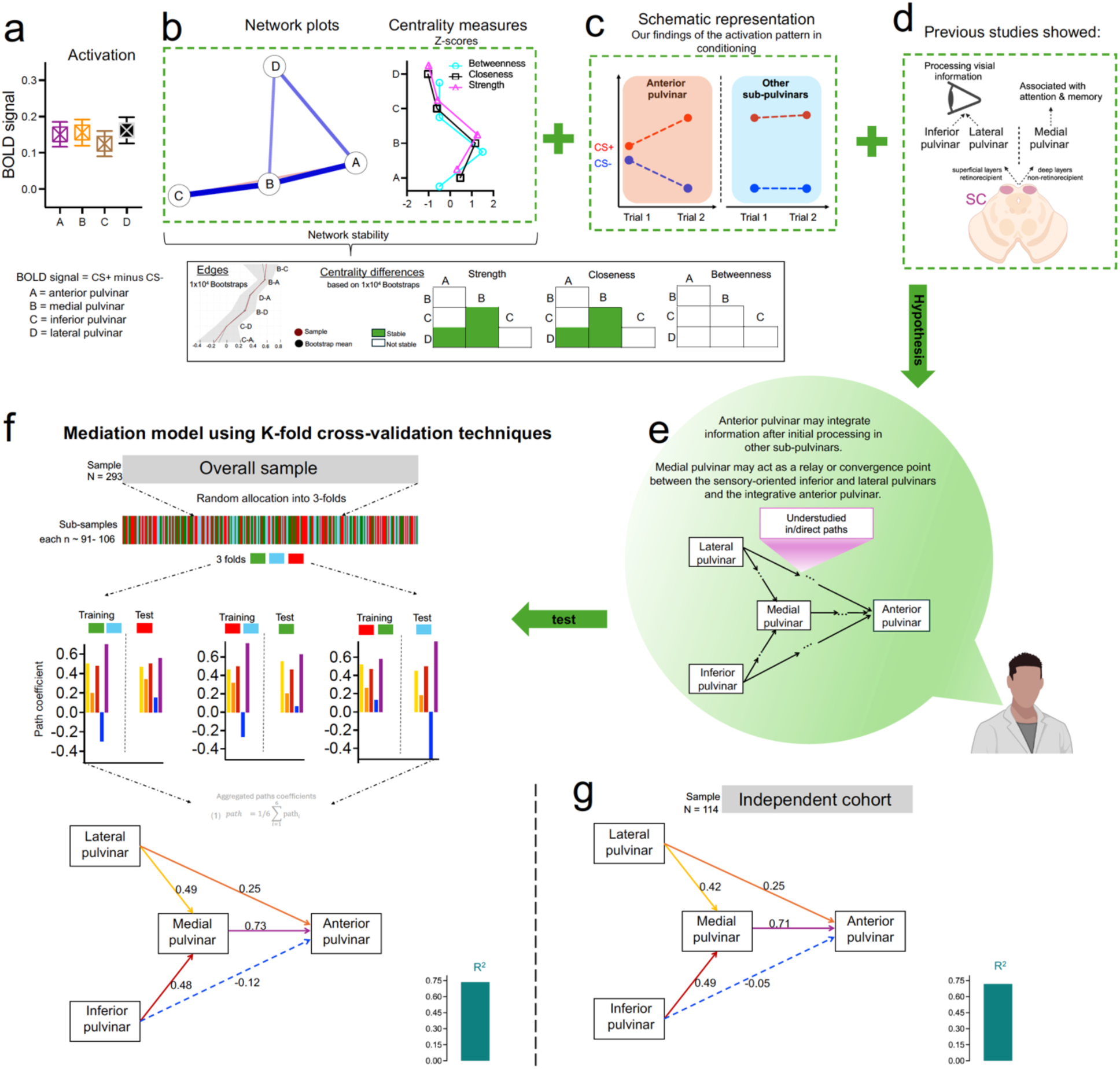
A data-driven approach to characterizing the functional organization of pulvinar divisions during conditioning. **(a)** Means ± SE of activation differences between pulvinar divisions. **(b)** Network analysis identified the medial pulvinar as a central node within the observed functional network, exhibiting the highest centrality measures. **(c)** Schematic visualization of activation patterns observed across pulvinar divisions. **(d)** Previous studies suggest that the inferior and lateral pulvinar are involved in processing basic visual information (Berman and Wurtz, 2008, 2010; Cortes et al., 2024), whereas the medial pulvinar has been associated with higher-order functions, including attention and working memory (Homman-Ludiye and Bourne, 2019). **(e)** Based on panels **b-d**, we propose a computational model in which medial pulvinar activation statistically accounts for the associations among activation in the other pulvinar divisions. **(f)** Statistical mediation analysis evaluating the proposed computational model. **(g)** Replication of the mediation model in an independent sample. This independent cohort (*N* = 114) was collected as part of a separate research project that included the same Day 1 fear-conditioning paradigm used in the present study. The cohort was used to assess whether the mediation findings could be replicated independently. Dashed paths in panels **f** and **g** represent statistically unstable paths, whereas solid paths indicate stable paths. **Note:** The functional relationships among pulvinar divisions during threat learning should be interpreted as computational dependencies derived from statistical associations rather than evidence of direct anatomical connectivity. These statistical associations may reflect indirect interactions involving broader corticothalamic and thalamocortical circuits (e.g., via visual cortical regions). Determining the anatomical mechanisms underlying these functional associations will require future studies. Supplementary analyses examining within-pulvinar functional associations while controlling for the effects of the neighboring thalamic nuclei included in this study (MD and LGN) are presented in **Supplementary Fig. 4**. These analyses provide an additional assessment of the specificity of the observed functional associations but do not distinguish direct inter-nuclear interactions from indirect coupling through corticothalamic and thalamocortical pathways. *p<0.05, **p<0.01, ***p<0.001. SE, standard error; SC, superior colliculus. Display items in panels **d** and **e** were created using BioRender (BioRender.com/1m4j5bz).

These network properties and trial-dependent activation patterns observed in the trial-wise analyses (overview in **Fig. 3c** and details in **Fig. 1b**) led us to propose a possible functional framework describing the statistical relationships among pulvinar divisions. Specifically, the increased activation induced by CS+ during trial 1 in the medial (*p_FDR_ = 0.026*), inferior (*p_FDR_ = 0.008*), and lateral pulvinar (*p_FDR_ = 0.01*) suggests that these divisions process CS+ information before the anterior pulvinar, which exhibited a delayed BOLD response starting in trial 2 only (*p_FDR_ = 0.44* in trial 1, while *p_FDR_ = 0.000001* in trial 2). The medial pulvinar statistically mediates the functional relationships within the pulvinar network and is associated with elevated centrality values. Together, these findings are consistent with a possible hub-like functional role of the medial pulvinar, consistent with higher-order integration of CS+ information, relative to the more sensory-related functional profiles in the inferior and lateral pulvinar. This possible hierarchical organization is consistent with previous evidence showing that the inferior and lateral pulvinar are associated with processing basic sensory information (Berman and Wurtz, 2010, 2008; Cortes et al., 2024), while the medial pulvinar is implicated in higher-order processing, including attention and working memory (Homman-Ludiye and Bourne, 2019) (**Fig. 3d**).

This data-driven analyses provide a possible new perspective on the functional specialization of pulvinar divisions during associative threat learning. Based on these observations, we hypothesized a computational model in which activation in the medial pulvinar statistically mediates the functional relationships between the inferior and lateral pulvinar and the anterior pulvinar (**Fig. 3e**). To evaluate this hypothesis, we performed a mediation analysis and assessed the robustness of the model using k-fold cross-validation. The sample (*N* = 293) was randomly divided into three groups of 91, 96, and 106 subjects. Each sub-sample was used to test the model while the remaining sub-samples were used to assess the stability of the parameter estimates. This resulted in six iterations of the mediation model. The 95% CI for each path coefficient were estimated using 10,000 bootstrapping replications. This choice was made to achieve greater precision and stability in the CI estimates. Finally, we applied the mediation analysis to an independent cohort (*N* = 114) collected as part of a separate research project using the same Day 1 fear-conditioning paradigm. The mediation findings were largely replicated in this independent cohort (**Fig. 3f-3g**). The results in **Fig. 3f-g** supported the proposed statistical mediation model, revealing significant indirect associations between activation in the lateral and inferior pulvinar divisions and activation in the anterior pulvinar that were statistically accounted for by activation in the medial pulvinar (*both p-values < 0.05*). Detailed statistical parameters are provided in **Supplementary Tables 3**.

### Pulvinar divisions and MD show engagement during extinction learning and context-dependent threat memory processing

After our comprehensive examination of the thalamic contribution to threat conditioning, we moved forward to examine the neural contributions of the same regions to extinction learning. During this phase, at the block level, we found higher activation in response to CS+E than CS- across all pulvinar divisions (*t_(319)_ = 2.02–2.85, p = 0.005–0.0.044, d = 0.11–0.15*) and MD (*t_(319)_ = 3.04, p = 0.003, d = 0.17)*. The trial-wise analysis indicated that these differences were primarily driven by the first two trials in the pulvinar; across pulvinar divisions, all second-trial comparisons survived FDR correction (*p_FDR_ < 0.05*, **Fig. 4b**), whereas the MD showed no significant effects at the trial-wise level (*all p_FDR_ > 0.05*, **Fig. 4c**). Thus, these findings indicate that pulvinar divisions and MD differed in the temporal pattern of CS+ versus CS− responses during extinction, with pulvinar differences observed at both trial and block levels, whereas MD differences were observed only at the block level. Detailed statistical parameters are provided in Supplementary Tables 4-5.

**Fig. 4.**
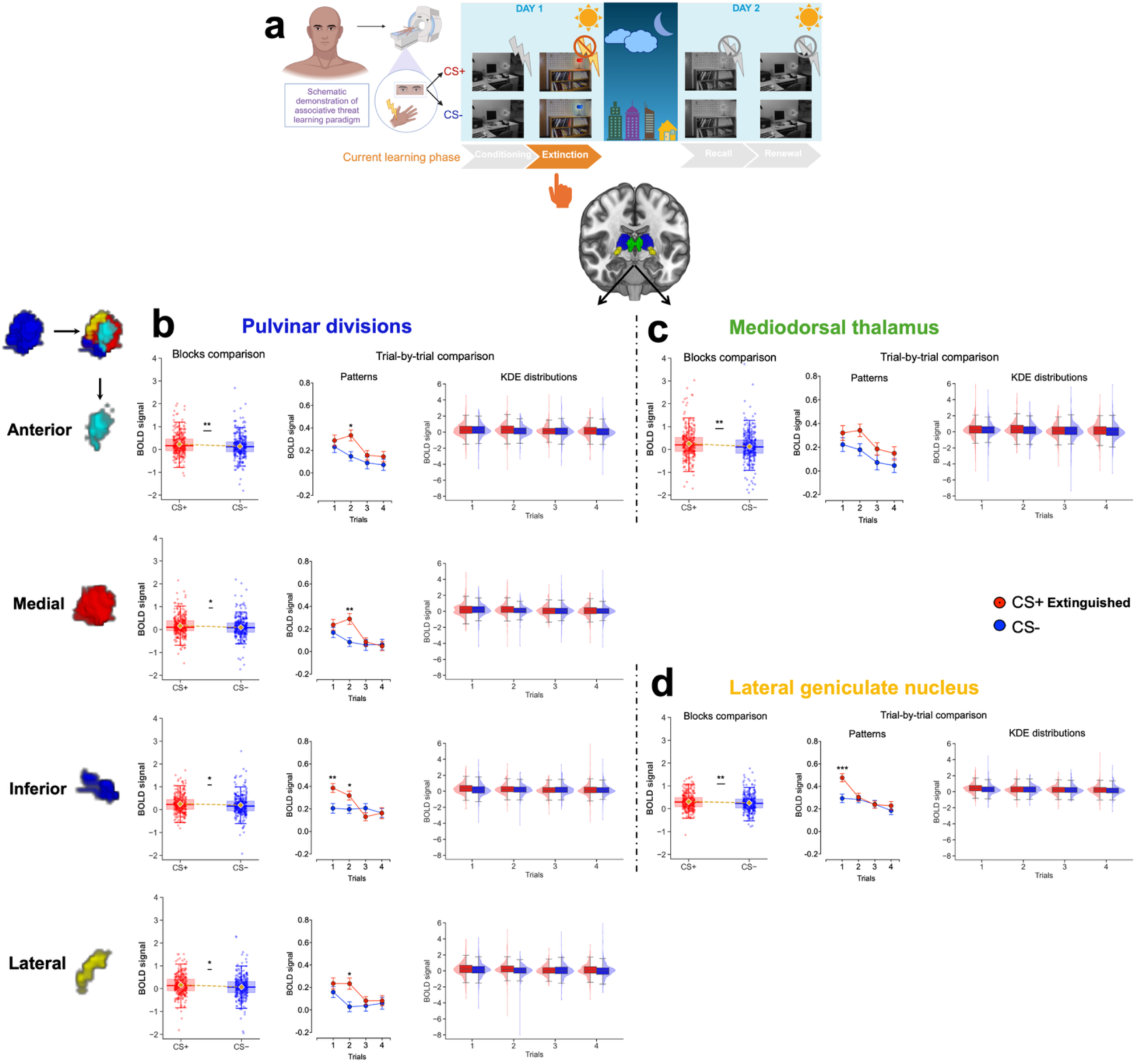
Neural representation during extinction learning in pulvinar divisions, MD, and LGN. (a) Human extinction learning paradigm during fMRI. (b-d) Boxplots with individual data points at the block level (gold diamonds indicate the means), and trial-wise activation shown as means ± SE and kernel density estimates (KDEs) for responses to CS+E and CS- in the pulvinar divisions (b), MD (c), and LGN (d). Block-level comparisons were assessed using paired *t*-tests, while trial-level effects were examined using a 2 × 2 repeated-measures ANOVA, followed by post hoc comparisons between CS+ and CS− across the four trials. Multiple comparisons were controlled using false discovery rate (FDR) correction. Extinction sample size: *n* = 320. Detailed statistical parameters are provided in **Supplementary Tables 4–5**. *p < 0.05, **p < 0.01, ***p < 0.001. SE, standard error; KDE, kernel density estimates; MD, mediodorsal thalamus; LGN, lateral geniculate nucleus; CS+E, extinguished conditioned stimulus that no longer predicts shock; CS−, conditioned stimulus predicting no shock. Display items in panel (a) were created using BioRender (BioRender.com/iori2z1).

During extinction recall, the anterior pulvinar and MD exhibited an increased activation in response to CS+E compared to CS- during the first two trials (Anterior pulvinar: at the block level *t*_(*411)*_ *= 2.49, p* = *0.013, d = 0.12, p_FDR_ = 0.026* in trial 1 and *p_FDR_ = 0.026* in trial 2; while *p_FDR_ > 0.05* in trial 3 and 4. MD: at the block level *t_(411)_ = 4.3, p = 0.00002, d* = *0.21, p_FDR_ = 0.000002* in trial 1 and *p_FDR_ = 0.008* in trial 2 while *p_FDR_ > 0.05* in trial 3 and 4; **Fig. 5b-c**). The medial and inferior pulvinar maintained the BOLD signal to CS+E in the second trial (both *p_FDR_* in trial 2 *< 0.01* while *p_FDR_* in other trials *> 0.05*). We observed no activation differences in response to CS+E vs. CS- in the lateral pulvinar (*p-values* at the block and trial-wise level *> 0.05*). See **Fig. 5b-c**, and Detailed statistical parameters in **Supplementary Tables 6-7**. These findings are consistent with engagement of most pulvinar divisions and the MD during the processing of threat-related memories under safe contextual cues and may reflect processes associated with the retrieval suppression of the extinguished threat.

**Fig. 5.**
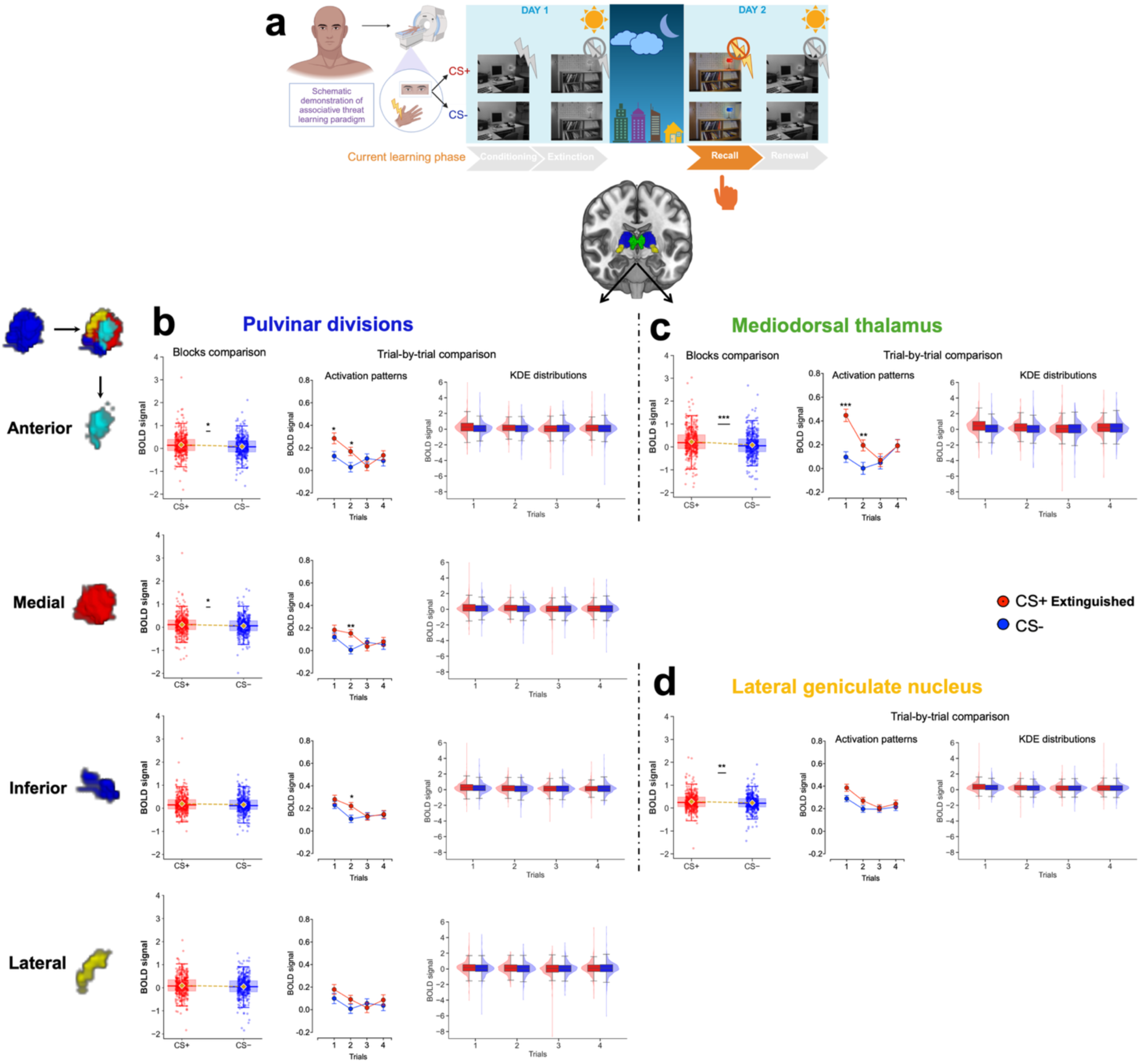
Neural representation during extinction recall in pulvinar divisions, MD, and LGN. (a) Human extinction recall paradigm in the fMRI conducted within safe contextual cues. (b-d) Boxplots with individual data points at the block level (gold diamonds indicate the means), and trial-wise activation shown as means ± SE and kernel density estimates (KDEs) for responses to CS+E and CS- in the pulvinar divisions (b), MD (c), and LGN (d). Block-level comparisons were assessed using paired *t*-tests, while trial-level effects were examined using a 2 × 2 repeated-measures ANOVA, followed by post hoc comparisons between CS+ and CS− across the four trials. Multiple comparisons were controlled using false discovery rate (FDR) correction. Extinction recall sample size: *n* = 412. Detailed statistical parameters are provided in **Supplementary Tables 6–7.** *p < 0.05, **p < 0.01, ***p < 0.001. SE, standard error; KDE, kernel density estimates; MD, mediodorsal thalamus; LGN, lateral geniculate nucleus; CS+E, extinguished conditioned stimulus that no longer predicts shock; CS−, conditioned stimulus predicting no shock. Display items in panel (a) were created using BioRender (BioRender.com/imzbkmn).

The threat renewal phase, in which participants were exposed to contextual cues similar to those used during threat conditioning, was associated with increased activation to CS+E on the first trial in the anterior pulvinar (*p_FDR_ = 0.0001*), lateral pulvinar (*p_FDR_ = 0.008*), and MD (*p_FDR_ = 0.0003*). No significant activation was observed in the medial or inferior pulvinar at the trial level (all *p_FDR_ > 0.05* across trials; **Fig. 6b–c**; Detailed statistical parameters are provided in **Supplementary Tables 8-9**.

**Fig. 6.**
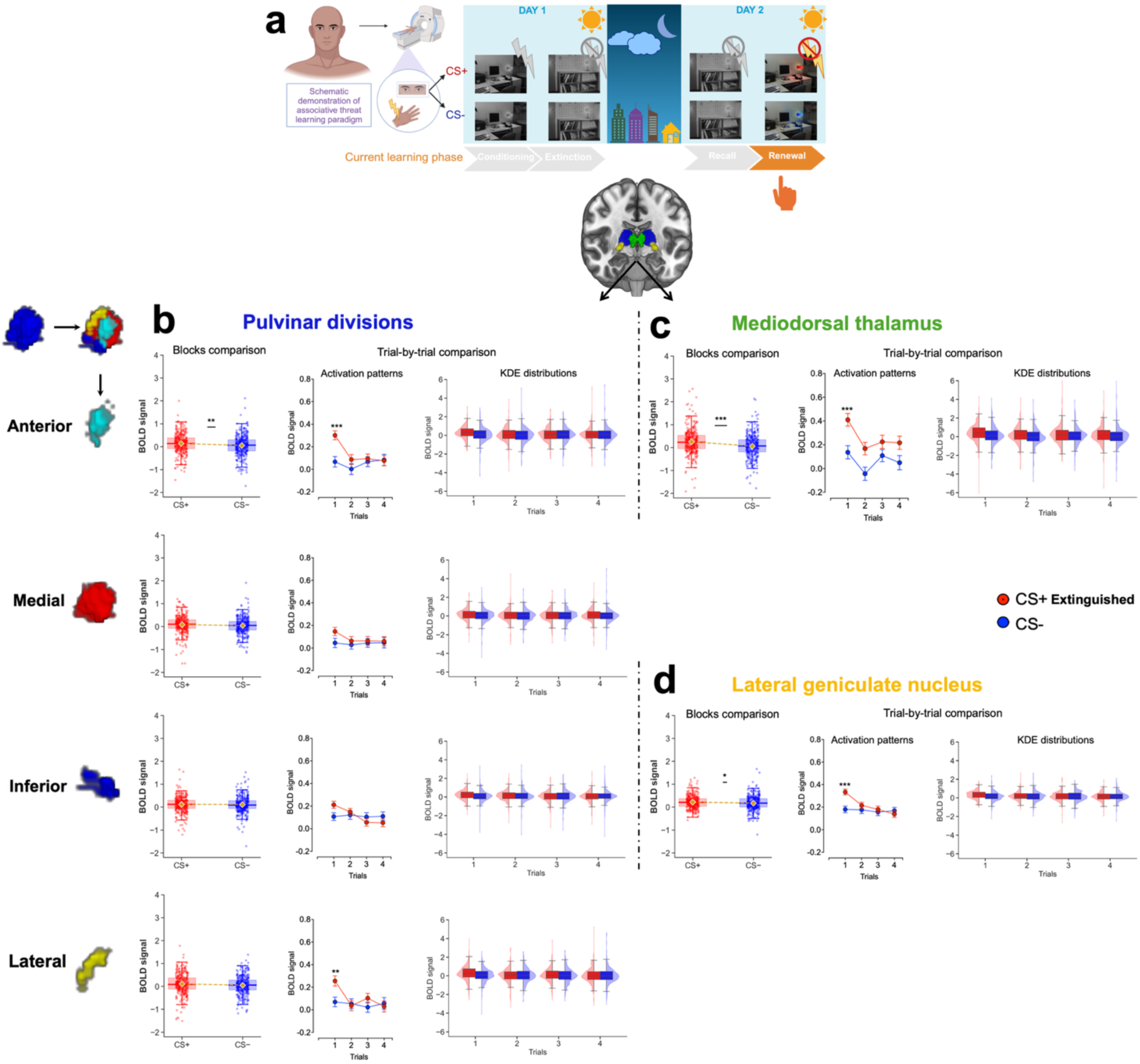
Neural representation during threat renewal in pulvinar divisions, MD, and LGN. (a) Human threat renewal paradigm in the fMRI conducted within threat contextual cues in the original context where fear conditioning occurred. (b-d) Boxplots with individual data points at the block level (gold diamonds indicate the means), and trial-wise activation shown as means ± SE and kernel density estimates (KDEs) for responses to CS+E and CS- in the pulvinar divisions (b), MD (c), and LGN (d). Block-level comparisons were assessed using paired *t*-tests, while trial-level effects were examined using a 2 × 2 repeated-measures ANOVA, followed by post hoc comparisons between CS+ and CS− across the four trials. Multiple comparisons were controlled using false discovery rate (FDR) correction. Threat renewal sample size: *n* = 318. Detailed statistical parameters are provided in **Supplementary Tables 8–9.** *p < 0.05, **p < 0.01, ***p < 0.001. SE, standard error; KDE, kernel density estimates; MD, mediodorsal thalamus; LGN, lateral geniculate nucleus; CS+E, extinguished conditioned stimulus that no longer predicts shock; CS−, conditioned stimulus predicting no shock. Display items in panel (a) were created using BioRender (BioRender.com/r9bhqjs).

### LGN activation patterns during threat learning seem to be consistent with feedforward processing

At the block level, activation was greater for CS+ than CS− within each phase separately: conditioning (*t_(292)_ = 3.35, p = 0.0009, d = 0.19*), extinction (*t_(319)_ = 2.66, p = 0.008, d = 0.14*), recall (*t_(411)_ = 2.61, p = 0.009, d = 0.12*), and renewal (*t_(317)_ = 2.41, p = 0.016, d = 0.13*). These differences were primarily driven by the first trial (*p_FDR_ < 0.001* in trial 1 across phases, except recall: *p_FDR_ = 0.056*), during which CS+ elicited increased BOLD responses that declined over subsequent trials. See **Figs. 1d**, **4d–6d**; Detailed statistical parameters are provided in **Supplementary Tables 2, 5, 7, and 9**. Collectively, these findings suggest that the LGN exhibited early trial-dependent responses within each learning phase separately, consistent with its proposed role in feedforward visual processing.

### Thalamic connectivity during threat learning and memory

We tested the relationships between each thalamic nucleus and core brain regions within the ‘fear circuit’ (Herry et al., 2010; Maren and Quirk, 2004; Milad et al., 2014; Milad and Quirk, 2012; Tovote et al., 2015) by seeding the nuclei to target the amygdala, hippocampus, ventromedial prefrontal cortex (vmPFC), subgenual anterior cingulate cortex (sgACC), and dorsal anterior cingulate cortex (dACC). During conditioning, the anterior pulvinar showed significant CS+ > CS− connectivity with the amygdala (*t_(292)_ = 2.46, p_FDR_ = 0.02, d = 0.14*), vmPFC (*t_(292)_ = 2.60, p_FDR_ = 0.02, d = 0.15*), and hippocampus (*t_(292)_ = 2.42, p_FDR_ = 0.02, d = 0.14*), but not with the dACC (*t_(292)_ = -0.95, p_FDR_ = 0.42, d = -0.05*) or sgACC (*t_(292)_ = 0.79, p_FDR_ = 0.42, d = 0.04*), **Fig. 7a**.

**Fig. 7.**
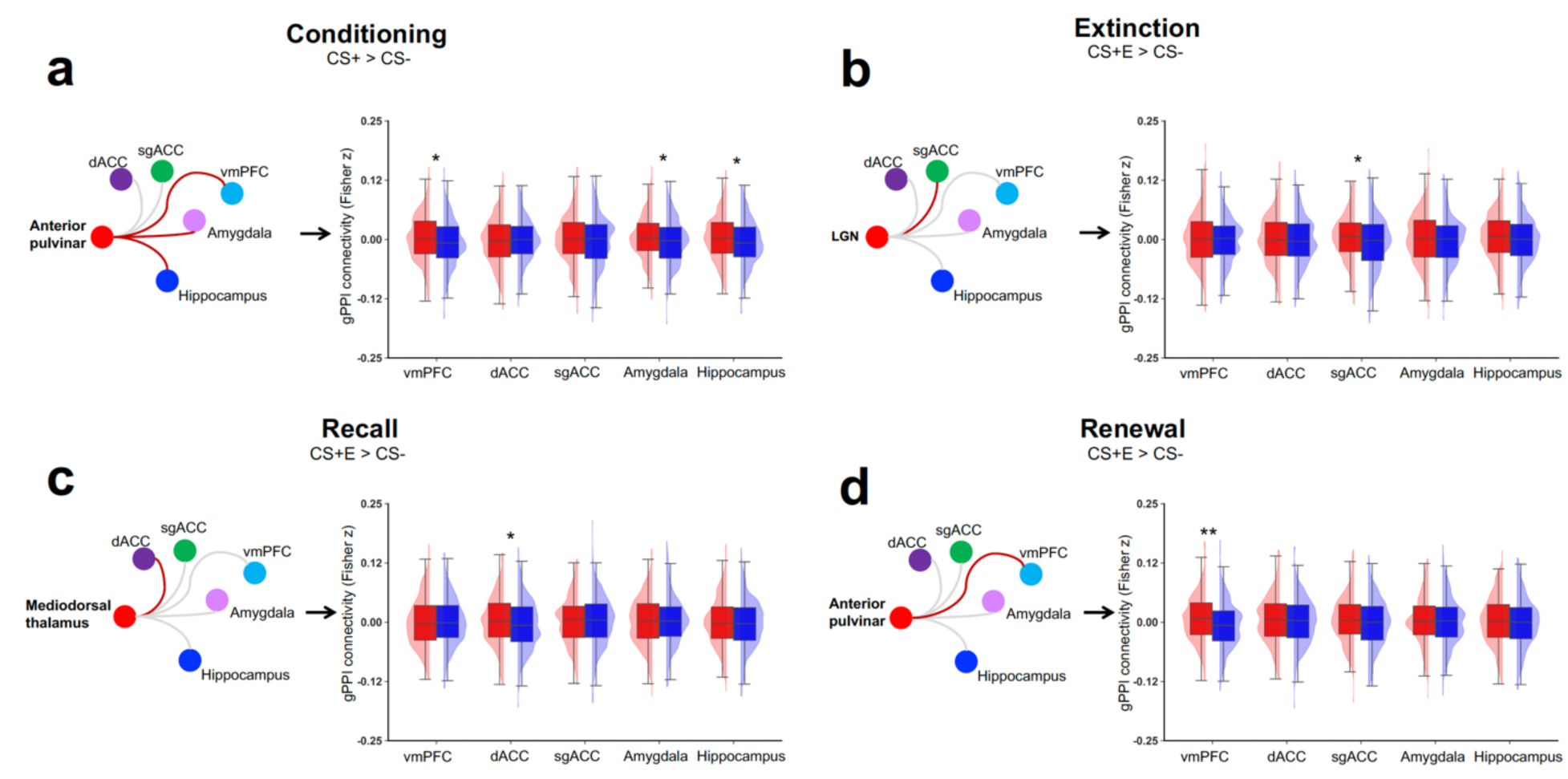
Task-related thalamic connectivity with threat-related brain regions. (a–d) Thalamic seed-to-ROI connectivity with the vmPFC, sgACC, dACC, amygdala, and hippocampus during conditioning (*n = 293*), extinction (*n = 320*), recall (*n = 412*), and renewal (*n = 318*). Red lines indicate significant positive connectivity for CS+ vs. CS− following FDR correction across regions (*p_FDR_ < 0.05*); gray lines indicate non-significant effects. For each panel, boxplots and kernel density estimates show the distribution of connectivity values for CS+ and CS−. All thalamic regions of interest—including pulvinar divisions (anterior, inferior, lateral, medial), LGN, and MD—were analyzed using identical statistical procedures. To maintain figure clarity and focus on effects relevant to the main conclusions, we display only seeds showing at least one significant CS+ vs. CS− differences after FDR correction. Non-significant seeds did not show systematic condition-related differences. *p < 0.05, **p < 0.01. MD, mediodorsal thalamus; LGN, lateral geniculate nucleus; vmPFC, ventromedial prefrontal cortex; sgACC, subgenual anterior cingulate cortex; dACC, dorsal anterior cingulate cortex; CS+, conditioned stimulus predicting shock; CS−, conditioned stimulus predicting no shock. gPPI; generalized psychophysiological interaction; connectivity values are Fisher z–transformed correlation coefficients.

Additionally, to assess the robustness of the connectivity differences between CS+ and CS−, we performed nonparametric bootstrapping (10,000 resamples) and estimated the bias-corrected and accelerated (BCa) CI for the mean paired differences. The BCa intervals excluded zero for the amygdala (BCa 95% CI [0.002, 0.015]), vmPFC (BCa 95% CI [0.002, 0.017]), and hippocampus (BCa 95% CI [0.0016, 0.016]), but included zero for the dACC (BCa 95% CI [-0.01, 0.003]) and sgACC (BCa 95% CI [-0.005, 0.011]). The connectivity with the amygdala may be consistent with the involvement of this circuit in processing the emotional valence of newly learned CS-US associations, while the engagement of the hippocampus may reflect the involvement of contextual information during CS+ processing. The vmPFC connectivity may reflect involvement of prefrontal processes related to evaluation and regulation of threat-related information.

During extinction learning, the LGN showed increased CS+E > CS− connectivity with the sgACC (*t_(319)_ = 2.60, p_FDR_ = 0.04, d = 0.14*). No significant differences were observed with other cortical regions, including the vmPFC (*t_(319)_ = 1.13, p_FDR_ = 0.32, d = 0.06*) and dACC (*t_(319)_ = 0.41, p_FDR_ = 0.68, d = 0.02*), or with limbic regions, including the amygdala (*t_(319)_ = 1.19, p_FDR_ = 0.32, d = 0.06*) and hippocampus (*t_(319)_ = 1.34, p_FDR_ = 0.32, d = 0.07*), **Fig. 7b**.

Additionally, to assess the robustness of the connectivity differences between CS+E and CS−, we performed nonparametric bootstrapping analysis (10,000 resamples). The BCa interval for the mean differences excluded zero for the LGN–sgACC connectivity (BCa 95% CI [0.002, 0.017]), but included zero for all other regions: vmPFC (BCa 95% CI [−0.003, 0.011]), dACC (BCa 95% CI [−0.005, 0.009]), amygdala (BCa 95% CI [−0.002, 0.012]), and hippocampus (BCa 95% CI [−0.002, 0.012]).

Interestingly, within each phase separately, the findings suggest that thalamic connectivity patterns vary with contextual conditions during extinction recall and threat renewal. Specifically, the MD and anterior pulvinar showed distinct CS+E vs. CS− connectivity patterns during these phases, respectively. During extinction recall, which occurs in the safe context, we observed increased connectivity to CS+E > CS- between the MD and dACC (*t_(411)_ = 2.60, p_FDR_ = 0.04, d = 0.12*) but not with other regions: amygdala (*t_(411)_ = 0.038, p_FDR_ = 0.97, d = 0.002*), hippocampus (*t_(411)_ = 1.2, p_FDR_ = 0.57, d = 0.059*), sgACC (*t_(411)_ = 0.18, p_FDR_ = 0.97, d = 0.009*), and vmPFC (*t_(411)_ = -0.49, p_FDR_ = 0.97, d = - 0.02*), **Fig. 7c**.

These findings were robust to nonparametric bootstrapping (10,000 resamples): the BCa 95% confidence interval for the mean connectivity difference excluded zero for the MD–dACC (BCa 95% CI [0.002, 0.015]), but included zero for the amygdala (BCa 95% CI [−0.006, 0.006]), hippocampus (BCa 95% CI [−0.002, 0.010]), sgACC (BCa 95% CI [−0.006, 0.007]), and vmPFC (BCa 95% CI [−0.008, 0.005]).

During threat renewal, we observed increased CS+E > CS− connectivity between the anterior pulvinar and the vmPFC (*t_(317)_ = 3.47, p_FDR_ = 0.002, d = 0.19*). In contrast, no significant CS+E > CS− connectivity differences were observed between the anterior pulvinar and other regions, including the amygdala (*t_(317)_ = 0.59, p_FDR_ = 0.55, d = 0.03*), hippocampus (*t_(317)_ = 1.09, p_FDR_ = 0.44, d = 0.06*), sgACC (*t_(317)_ = 1.90, p_FDR_ = 0.14, d = 0.10*), and dACC (*t_(317)_ = 0.92, p_FDR_ = 0.44, d = 0.05*), **Fig. 7d**.

Nonparametric bootstrapping (10,000 resamples) supported the robustness of these results. The BCa 95% confidence interval for the mean difference excluded zero only for the anterior pulvinar –vmPFC connectivity (BCa 95% CI [0.005, 0.02]), whereas intervals for the amygdala (BCa 95% CI [-0.005, 0.01]), hippocampus (BCa 95% CI [-0.003, 0.012]), sgACC (BCa 95% CI [-0.0002, 0.015]), and dACC (BCa 95% CI [-0.003, 0.01]) all included zero.

## Discussion

We examined the neural representation of associative threat learning within the pulvinar divisions, LGN, and MD, providing new insights into thalamic involvement in this adaptive behavior in humans. We identified distinct patterns of thalamic activation during threat learning and memory. The anterior pulvinar and MD exhibited parallel activation patterns consistent with associative learning, raising a possibility that these nuclei may be involved in automatic survival responses and deliberate threat processing. The medial pulvinar may statistically mediate the functional relationships between the inferior and lateral divisions and the anterior pulvinar, consistent with a hierarchical computational model. Both the pulvinar and MD showed activation patterns that were consistent with salience processing of threat-related memories when threat-related cues were encountered during extinction recall and threat renewal (considered separately in each phase). The LGN exhibited activation patterns that may be consistent with feedforward visual processing, anticipating upcoming visual stimuli throughout threat learning. We integrated these insights into schematic models underlying the emotional and behavioral expression of threat learning and memory, providing a possible neural framework for studying related human behaviors (**Fig. 8a-b**).

**Fig 8.**
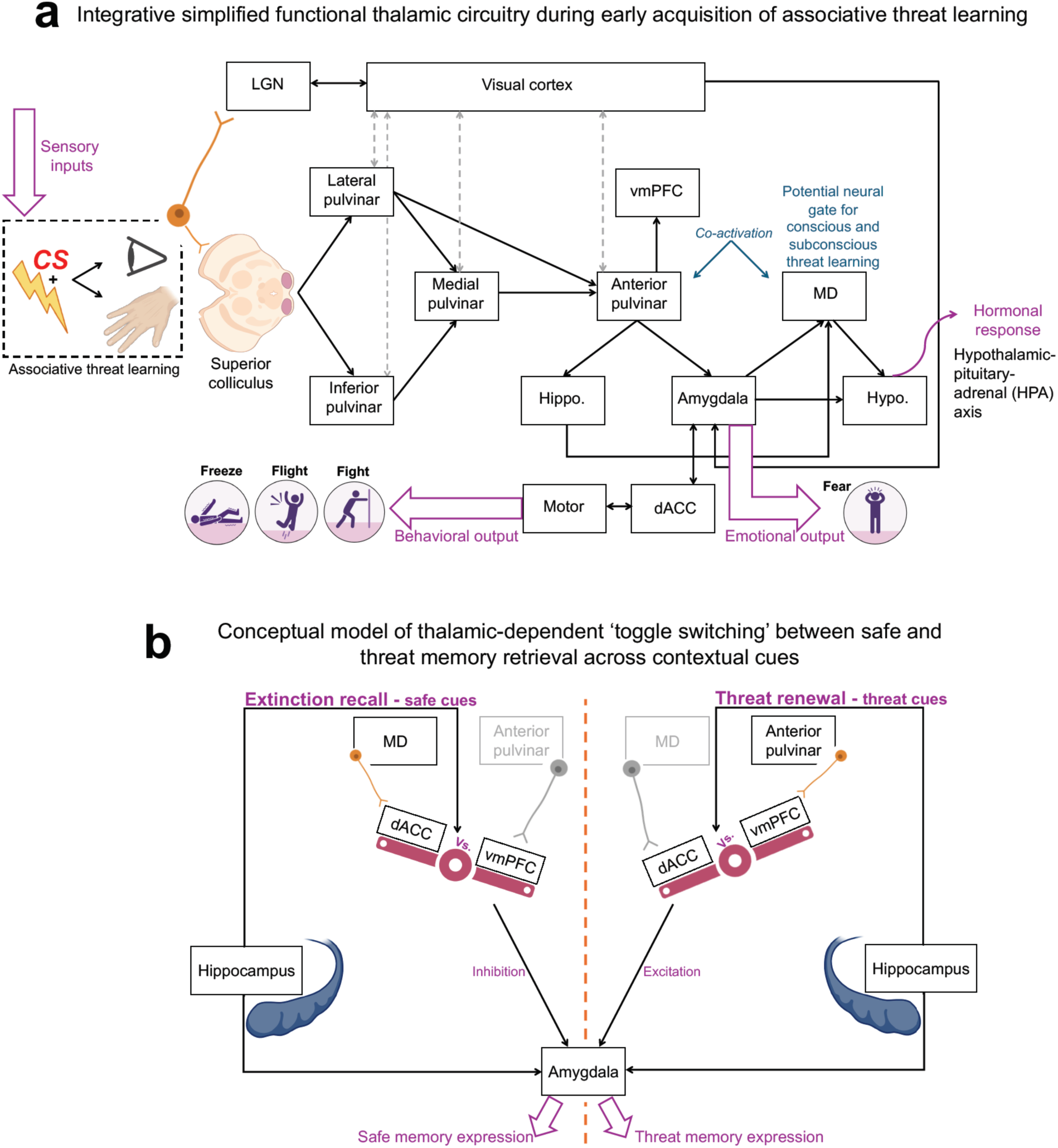
Proposed conceptual neurobehavioral models of thalamic involvement in associative threat learning and memory. (a) Conceptual schematic illustration of thalamic circuitry during the acquisition of associative threat learning, highlighting interactions between pulvinar divisions, MD, and LGN with key brain regions implicated in threat-related processing. (b) Conceptual model illustrating a possible thalamic-dependent “toggle switch” underlying context-dependent safety and threat memory expression. The observed MD-dACC and anterior pulvinar-vmPFC connectivity patterns are consistent with possible preferential engagement of prefrontal circuits during extinction recall and threat renewal, respectively. This framework represents a theoretical integration of the present findings rather than evidence of causal neural mechanisms. It is based on the findings observed within each phase separately and should not be interpreted as a direct comparison between phases. **Note (panel a):** Pulvinar-cortical pathways, as well as sensory input pathways (e.g., visual inputs via the retina/LGN and nociceptive inputs via the spinothalamic tract), are not explicitly shown. The pulvinar-cortical pathways and major sensory input pathways are recognized anatomically but were not specified due to their heterogeneity and incomplete characterization at the level of pulvinar subnuclei. Their lack of specification should not be interpreted as a lack of anatomical or functional relevance. The bidirectional dashed gray arrows between the pulvinar and visual cortex schematically represent corticothalamic and thalamocortical pathways. Their precise organization and contribution to individual pulvinar subnuclei remain incompletely characterized and were not examined in the present study. MD: Mediodorsal thalamus. LGN: Lateral geniculate nucleus. vmPFC = Ventromedial prefrontal cortex. dACC = Dorsal anterior cingulate cortex. Hypo.: Hypothalamus. Hippo.: Hippocampus. Created using BioRender (BioRender.com/5pdf54t).

The anterior pulvinar and MD co-activation patterns may reflect a parallel contribution to threat learning. The similar activation in response to CS+ and CS- at the first trial, followed by a heightened activation specific to CS+ in the next trial, is likely consistent with the acquisition of the CS-US association. The responses’ similarity to both types of CS during the first trial suggests a possible equivalent initial emotional valence. The gradual increase in the activation, specifically to CS+ by the end of trial 1, is consistent with a shift in the emotional valence of CS+, likely indicating rapid associative learning. Together, these findings are consistent with the possibility that the anterior pulvinar and MD contribute in parallel to associative threat learning, extending the two-system model proposed by LeDoux and Pine (LeDoux and Pine, 2016). Briefly, fear processing, according to this model, involves two distinct pathways: a subcortical route for rapid threat detection and species-specific defense reactions and a cortical route for deliberate threat evaluation and conscious fear experience. Anterior pulvinar activation may be consistent with rapid automatic processes proposed for the subcortical pathway. This aligns with previous reports that demonstrated the role of the pulvinar-amygdala pathway in encoding negative emotions in humans (Bridge et al., 2015; Koller et al., 2019; Kragel et al., 2021; McFadyen et al., 2019; Rafal et al., 2015). Indeed, the increased connectivity that we observed with the amygdala, hippocampus, and vmPFC during threat conditioning is consistent with this interpretation. Previous work suggests that the amygdala contributes to tagging a negative emotional valence to CS+ and triggering fight, flight, or freeze reactions (Adolphs et al., 1994; Costa et al., 2022; Fanselow and LeDoux, 1999; Phelps and LeDoux, 2005; Wen et al., 2024, 2022). The observed hippocampal connectivity may be consistent with contextual encoding of the environmental characteristics of the aversive event (Maren, 2001; Maren et al., 2013). The observed vmPFC connectivity may reflect processes related to salience encoding, future threat evaluation (Battaglia et al., 2020), decision-making, and emotion regulation (Milad and Quirk, 2012, 2002; Nejati et al., 2021).

On the other hand, the MD activation pattern may be more consistent with the cortical pathway proposed in the two-system model, which has been associated with conscious and deliberate threat processing. This interpretation is supported by our finding that MD activation in response to CS+ was greater than that of the anterior pulvinar during threat learning, consistent with the possibility that these nuclei contribute differently within the two-system framework. Additionally, this interpretation is supported by substantial evidence pointing to well-established anatomical pathways connecting the MD with the PFC and demonstrating its role in decision-making and learning (Behrens et al., 2003; Hwang et al., 2020; Li et al., 2022; Wolff and Halassa, 2024). This is along with reports that underscored the contribution of the MD-cortical loops to supporting a conscious experience in humans (Griffiths et al., 2022; Whyte et al., 2024). Together, these findings align with the framework of the two-system model of threat processing, raising the possibility that the anterior pulvinar and MD possible role in the acquisition of the CS-US association and suggesting that these regions might be involved in a parallel contribution to conscious vs. unconscious aspects of threat learning proposed by LeDoux and Pine (LeDoux and Pine, 2016), **Fig. 8a**.

These findings are consistent with rapid initial acquisition of the predictive value of the CS+ is thought to be pronounced during the first two trials. The attenuated CS+ vs. CS− differentiation on the third trial specifically in the pulvinar may reflect a decreased requirement for differential thalamic engagement once the initial association has been acquired. Notably, because the MD sustained the BOLD response to the CS+ in the third trial which may indicate involvement of this nucleus in continued processing or maintenance of the learned association. This aligns with the well-established MD-PFC circuit involved in cognitive processes (Wolff and Halassa, 2024). Additionally, in a previous study using a similar paradigm, we observed sustained CS+ vs. CS− differentiation on the third trial in the nucleus reuniens, as well (Tuna et al., 2025). These findings suggest that trial-dependent learning dynamics may vary across thalamic nuclei rather than reflecting a uniform thalamic learning signal. Together, while our paradigm does not inherently distinguish between different stages of learning, such as early acquisition and stabilization, our findings are consistent with stronger associative learning–related engagement during the first two trials, with a reduced differential response by the third trial that may reflect the involvement of different neural processes.

We propose that the observed activation patterns across pulvinar divisions are consistent with a hierarchical computational organization associated with threat processing. Although the anatomical projections within pulvinar divisions are not well characterized, our functional data-driven approach suggests a possible critical role for the medial pulvinar as a hub region within the observed functional network. Specifically, our trial-wise analysis suggested that the medial, inferior, and lateral pulvinar showed earlier CS+ related activation than the anterior pulvinar. The anterior pulvinar exhibited activation in response to CS+ starting at trial 2, whereas activation in the other pulvinar divisions occurred as early as trial 1. Network modeling further identified the medial pulvinar as a potential central hub, likely facilitating communication with other divisions. Using statistical mediation analysis, we found that activation in the medial pulvinar statistically mediated the relationship between activation in the inferior/lateral pulvinar and activation in the anterior pulvinar. This statistical mediation is consistent with, but does not establish, a hierarchical computational organization. This proposed hierarchical model aligns with previous evidence suggesting that the inferior and lateral pulvinar are associated with processing basic sensory information (Berman and Wurtz, 2010, 2008; Bridge et al., 2015; Cortes et al., 2024). While the medial pulvinar, which receives projections from deep layers of the superior colliculus, supports more advanced functions such as attention and working memory (Bridge et al., 2015; Homman-Ludiye and Bourne, 2019). These findings seem to be indicative of novel insights into the functional specialization of pulvinar divisions during threat learning, consistent with proposed feedforward roles in the inferior and lateral pulvinar and high-order integration in the anterior pulvinar.

Yet, these intrapulvinar relationships should be understood as a functional and computational model, reflecting statistical associations among pulvinar divisions during threat learning, rather than as evidence of direct monosynaptic anatomical connections. Because detailed inter-nuclear anatomical connectivity within the pulvinar remains incompletely characterized, our analysis does not presuppose strong direct excitatory projections between subnuclei. Instead, our findings are intended to highlight candidate functional relationships within the pulvinar during conditioning, rather than to provide a definitive anatomical map.

The pulvinar divisions and the MD showed patterns consistent with engagement during extinction learning. In the pulvinar, responses to the CS+E remained relatively stable across the first two trials and subsequently attenuated; a pattern that may be consistent with rapid updating of threat value. In contrast, MD activation patterns were more consistent with sustained block-level responses across extinction, which may reflect sensitivity to broader task-related information. When examined separately, extinction recall and threat renewal each showed continued responses to the CS+E under their respective contextual conditions. This pattern may be consistent with retrieval of learned threat-related information during context-dependent memory processing. During extinction recall, this pattern may be consistent with retrieval suppression, whereas during threat renewal it may reflect continued processing of threat salience. Together, these findings suggest that the MD and pulvinar remain responsive to the extinguished CS+ across contextual conditions rather than exhibiting a static pattern of responding.

The classical Pavlovian model proposes that extinction learning forms a new safety memory, competing with the original threat memory acquired during conditioning (Bouton, 2002; Milad and Quirk, 2002). Contextual changes often favor either the safe or threat memory. Maren and colleagues (Maren et al., 2013; Maren and Quirk, 2004) described the hippocampal-prefrontal-amygdala model for contextual memory. The hippocampus projects to the basolateral amygdala, vmPFC, and dACC. Although the vmPFC supports safe memory recall by projecting to the intercalated cells (which inhibit the central nucleus of the amygdala), the dACC enhances threat memory renewal by projecting to the basolateral amygdala (which activates the central nucleus of this structure). Our findings suggest that the thalamic connectivity may be associated with the balance between safe and threat memory by modulating the competitive interaction between the dACC and vmPFC. The increased MD-dACC connectivity during recall may be consistent with involvement of MD-prefrontal interactions associated with safe memory expression. On the other hand, the anterior pulvinar-vmPFC connectivity during fear renewal may be associated with context-dependent recruitment of prefrontal circuits during threat memory expression, may be associated with context-dependent recruitment of prefrontal circuits during threat memory expression. The thalamic connectivity, thus, may be associated with a context-dependent functional balance between the dACC and vmPFC. This balance could represent a conceptual ‘toggle switch’ framework between safe and threat memory expression. We integrated this suggested flexible responding mechanism to changing environmental contexts in Maren’s circuit model of emotional memory, **Fig. 8b**.

LGN activation showed a similar trial-dependent pattern across the learning phases examined separately, as evidenced by our trial-wise results, which pointed to increased activation in response to CS+ that diminished immediately after the first trial. This distinct pattern may reflect anticipation of upcoming visual stimuli. Within each phase examined separately, LGN showed a similar pattern of CS+-related activation. This aligns with previous studies that highlighted the LGN’s engagement in selective attention and anticipation of visual stimuli (Mahoney and Schmidt, 2024; O’Connor et al., 2002; Saalmann and Kastner, 2009). The LGN, as a first-order nucleus (Cortes et al., 2024), has been proposed to primarily supports a feedforward visual processing, while the pulvinar, as part of the visual thalamus, has been associated with broader visual and cognitive functions (Cortes et al., 2024) including emotional valences of stimuli. This distinction likely underscores their joint but specialized contributions to adaptive threat responses.

### Limitations and Future Directions

Although our findings suggest distinct patterns of thalamic involvement during threat learning, fMRI cannot fully capture the complexity of thalamic organization. Pulvinar divisions, MD, and LGN each contain diverse neuronal subtypes and finer anatomical subdivisions that may serve distinct functions. Importantly, the absence of CS+ vs. CS− differences in BOLD activation should not be interpreted as a lack of stimulus-specific processing, as such distinctions may occur without detectable changes in overall BOLD activation. Future advances in higher-resolution imaging and human brain atlases may further improve anatomical and functional precision. The present analyses were performed in standardized MNI space to enable consistent group-level analyses and ROI definition across participants. Although this is a widely accepted approach in human neuroimaging, future studies may examine the generalizability of the present findings using complementary approaches, including analyses performed in native subject space. The reduced CS+ vs. CS− differentiation observed in the pulvinar subdivisions during the third trial may reflect a transition from the rapid associative learning processes engaged during the first two trials to a different stage of threat processing. However, the present experimental design does not allow us to determine the specific psychological or neural mechanisms underlying this change. Importantly, this attenuation was not observed uniformly across thalamic nuclei, suggesting that it does not reflect a general thalamic response but may instead indicate nucleus-specific functional dynamics. Future studies specifically designed to dissociate different stages of threat learning will be required to determine the functional significance of this trial-dependent effect. Additionally, since different sensory modalities preferentially engage distinct thalamic nuclei, the specific thalamic roles we investigated may not be consistent across different experimental designs, particularly in studies using auditory threat learning. The observed relationships among pulvinar divisions during conditioning are purely functional and do not distinguish direct inter-nuclear interactions from indirect coupling mediated by corticothalamic and thalamocortical pathways, including visual cortical regions. Thus, the proposed functional pulvinar model may reflect indirect corticothalamic and thalamocortical loops, currently undocumented inter-nuclear interactions, or a combination of these mechanisms. Finally, given the indirect nature of fMRI data and the absence of direct brain signal manipulations, our findings should not be interpreted as evidence of causality. Further research is needed to examine causal mechanisms underlying dynamic neural representations of threat learning within the thalamus.

This study’s insights raise critical future questions at the intersection between neuroscience, psychology, and mental health, particularly regarding the neural mechanisms underlying associative threat learning. First, the observed involvement of the anterior pulvinar and MD during the acquisition of the CS-US association motivates future work aimed at developing more precise brain interventions, focusing on refining maladaptive fear reactions. Future studies could explore the optimal parameters for applying non-invasive techniques to modulate activation within thalamic circuits during or immediately after fear conditioning. If supported by future causal studies, this approach may prove relevant for clinical populations and individuals exposed to high-risk environments, such as emergency medical staff, firefighters, and paramedics.

Second, differential patterns of MD-dACC and anterior pulvinar-vmPFC connectivity during safe and threatening memory suggest potential intervention targets for prioritizing one memory over the other. Future studies could test whether modulation of MD-dACC circuitry facilitates safe memories while engaging the anterior pulvinar-vmPFC circuits might reinforce fear memory pathways. Experimental designs in controlled laboratory settings could target these circuits to identify specific parameters for enhancing safe memories. If validated experimentally, this avenue holds therapeutic potential, particularly in conditions such as PTSD and phobias.

Third, the anterior pulvinar-MD relationships raise questions related to the wide range of human behaviors, focusing on understanding neural gateways between conscious and unconscious learning. Studying the dynamics of information flow across this pathway could deepen our understanding of how sensory and cognitive information transits between conscious and unconscious states and influence behavior. This research line may provide new insights into innovative learning methods and reveal neurocognitive mechanisms underlying the acquisition of new information in humans.

Together, our findings and emerging research directions underscore the importance of thalamic nuclei in associative threat learning and memory, offering possible avenues for theoretical advancements and clinical applications. As the primary neural gateway to the human brain, the thalamus is the first station for all sensory information except olfactory inputs. Future studies may determine whether targeted modulation of thalamic circuits can produce widespread functional changes across brain networks and influence behavior.

## Materials and Methods

### Participants

We analyzed fMRI data of 293 participants during fear conditioning, 320 during extinction, 412 during extinction recall, and 312 during threat renewal. Participants included individuals of both sexes, aged 18–70 years (mean ± SD = 32.17 ± 13.1 years). All participants were proficient in English, right-handed, and had normal or corrected-to-normal vision. The exclusion criteria included a history of seizures or significant head trauma, current substance abuse or dependence, metal implants, pregnancy, breastfeeding, or positive urine toxicology screen for drugs of abuse. We followed the latest version of Helsinki’s declaration, and all procedures were approved by the Partners HealthCare Institute Review Board of the Massachusetts General Hospital, Harvard Medical School. All subjects provided written informed consent before taking part in the study. Results from this dataset have been published elsewhere with a different focus (Marin et al., 2020; Wen et al., 2024, 2022). The current results are novel and have not been previously published.

### Experimental procedure

#### Threat learning

Participants underwent two-day sessions of a validated threat learning paradigm in the MRI scanner (**Figs. 1a, 4a-6a**); the experimental contexts consisted of visual scenes on a computer display. On day 1, participants underwent a Pavlovian conditioning phase in which a neutral stimulus in context A (e.g., red light in an office) was paired with a 500ms electric shock (US) with a partial reinforcement rate of 62.5%. Another neutral stimulus in context A (e.g., blue light in the same office) was also presented but never paired with the shock (CS-). On the same day, we also conducted extinction learning in context B (e.g., red and blue lights in a casebook background). The stimuli were repeatedly presented with the removal of the expected reinforcement (i.e., no shock).

On day 2, the participants underwent two phases of a memory test. The first is extinction recall, including presenting the extinguished CS+ and CS- within safe contextual cues (context B used during extinction learning). The second is threat renewal, in which the stimuli were presented within threat contextual cues (context A used in conditioning). Both extinction recall and threat renewal included no US reinforcement. Across phases of threat learning, the duration of each trial was 6s, and the inter-trial intervals with a fixation screen ranged between 12 to 18 s (15 s on average). Finally, to control for potential confounds related to the experimental design, we pseudorandomized and counterbalanced the order of CS+ and CS− across phases and between subjects.

#### Shock level

The shock intensity used in the fear learning paradigm was determined during a pre-experiment calibration. Electrical stimulation was delivered using a BIOPAC stimulation system with BIOPAC electrodes and conductive gel. The electrodes were attached to the palm of the participant’s right hand and positioned in close proximity without making contact with one another. The electric stimulation began at a low level (0.1 mA), gradually increasing in small increments. After each increment, participants verbally rated their discomfort. The procedure continued until the participant identified a level they described as “highly annoying but not painful.” This individualized intensity was then used for that participant throughout the experiment. For safety and ethical reasons, the maximum intensity was capped at 20 mA, and no participant received a shock above this limit.

#### Instructions to the participants

Each visual stimulus in our paradigm was first shown to participants for 6 seconds. This initial presentation served as habituation, allowing us to isolate the responses to genuinely new stimuli. Before the experiment began, participants were informed that they would see pictures illuminated with different colored lights, such as red or blue. During the experiment, some pictures might be paired with an electric shock, while others might not. Participants were instructed to pay attention to whether a specific color or pattern was associated with the shock. These instructions were adopted from previous studies in which our group developed this paradigm and found them highly effective for human learning. We therefore used the same approach in the current experiment. These instructions were provided throughout all phases of threat learning, and participants were informed that any shocks delivered would be at the same intensity determined on Day 1.

### MRI data acquisition and preprocessing

Two MRI settings were used to acquire the neuroimaging data. The first is a Trio 3T whole-body MRI scanner (Siemens Medical Systems, Iselin, New Jersey) using an 8-channel head coil. The functional data in this setting were acquired using a T2* weighted echo-planar pulse sequence with these parameters: TR = 3.0s, TE = 30 ms, slice number = 45, voxel size = 3 × 3 × 3 mm^3^. The second setting was also in the same scanner using a 32-channel head coil. The functional images were obtained using a T2* weighted echo-planar pulse sequence using TR = 2.56s TE = 30 ms, slice number = 48, voxel size = 3 × 3 × 3 mm^3^. The anatomical brain images were collected using a T1-weighted MP-RAGE pulse sequence, parcellated into 1 × 1 × 1 mm^3^ voxels. Elastic bands were affixed to the head coil device to reduce head motions.

Using the default pipeline in fMRIPrep, version 20.0.2 (Esteban et al., 2020, 2019), we preprocessed the data and applied correction of slice timing, realignment of the functional images, and coregistration. In addition, spatial normalization was performed with data normalized to Montreal Neurological Institute (MNI) space and resampled to a 2 × 2 × 2 mm³ voxel grid, followed by spatial smoothing with a 6-mm full-width at half-maximum Gaussian kernel.

### Activation analyses

We applied the least-squares-based generalized linear model (GLM) for each participant to estimate the BOLD response to CS+ and CS- using SPM 12. We estimated the beta values for each voxel during each learning phase in the paradigm. Overall, the model included 32 regressors for the CS+ and CS-, a regressor for the context, and a regressor for shock in conditioning but not in other phases. The GLM also included six head movement parameters (x, y, z directions, and rotations). This first-level analysis resulted in contrast maps that we used to estimate the variability of these maps across all subjects at the group-level analysis. We then used the contrast maps from the group-level analysis to extract the averaged values across the voxels within predefined masks of the pulvinar divisions, MD, and LGN. These outputs were used to compare the BOLD response during the first four CS+ and four CS- trials. We averaged the activation for each CS type across trials to obtain the block-level activation.

### Statistical tests

All statistical analyses were conducted using JASP versions 0.18.3 and 0.19.3 (JASP Team, 2024).

### Activation differences across threat-learning phases

For each threat-learning phase, block-level activation was defined as the average activation across trials. Differences in activation between CS+ and CS− conditions within each thalamic nucleus were assessed using paired-samples t-tests.

To examine trial-wise activation dynamics, we conducted 2 × 2 RM-ANOVAs with condition (CS+, CS−) and Trial as within-subject factors. Assumptions of sphericity were evaluated using Mauchly’s test, and Greenhouse–Geisser corrections were applied when violations were detected. Significant effects were followed by post hoc comparisons at the trial level, with false discovery rate (FDR) correction applied to control for multiple comparisons.

### Network analyses of within-pulvinar relationships during conditioning

Network analyses examined functional relationships between pulvinar divisions. Nodes corresponded to block-level CS+ minus CS− activation estimates for each pulvinar division, yielding four nodes (one per division). Networks were estimated using a Gaussian graphical model with EBICglasso (LASSO regularization) based on Pearson correlation matrices, with the EBIC tuning parameter set to γ = 0.5. Edge weights represent partial correlations.

Three centrality measures were computed on the estimated weighted partial-correlation network: node strength, defined as the sum of absolute edge weights connected to a node; closeness, defined as the inverse of the average shortest path length from a node to all other nodes; and betweenness, defined as the proportion of shortest paths between all node pairs that pass through a given node. Shortest paths were computed using inverse edge weights. Centrality indices were normalized. Network accuracy and centrality stability were assessed using nonparametric bootstrapping (10,000 iterations) to estimate CI for edge weights and centrality measures. All network analyses were conducted using the default settings unless otherwise specified, following Epskamp et al., (2018)

### Mediation analyses of within-pulvinar relationships during conditioning

Mediation models were estimated using the lavaan package (Rosseel, 2012) with maximum likelihood estimation. All variables were z-standardized prior to analysis. Block-level activation estimates from the inferior and lateral pulvinar were entered as predictors, activation in the medial pulvinar as the mediator, and activation in the anterior pulvinar as the outcome.

To assess the robustness and generalizability of the mediation effects, we conducted K-fold cross-validation (*K = 3*). The full sample (*N = 293*) was randomly partitioned into three non-overlapping sub-samples (*n = 91, 96,* and *106*). In each iteration, the mediation model was estimated in one sub-sample, and consistency of indirect effects was evaluated across the remaining sub-samples, yielding six cross-validation iterations.

Indirect effects were evaluated using BCa bootstrap CI based on 10,000 resamples, as recommended by Biesanz et al., (2010). An indirect effect was considered significant when the 95% confidence interval did not include zero and *p < 0.05*.

### Connectivity analyses

Functional connectivity was computed using the CONN toolbox (version 22.a) for MATLAB (Nieto-Castanon and Whitfield-Gabrieli, 2022; Whitfield-Gabrieli and Nieto-Castanon, 2012). Anatomical images were segmented into gray matter, white matter, and CSF. Functional images were preprocessed following CONN’s default pipeline, including realignment, slice-timing correction, and coregistration. Functional images were then denoised using a component-based noise correction approach (CompCor), including regression of estimated motion parameters, regression of identified outlier scans using artifact detection procedures (scrubbing), and regression of white matter and CSF signals. The denoised BOLD time series were subsequently band-pass filtered between 0.008 and 0.09 Hz.

Following these denoising procedures – including regression of motion parameters, artifact-related outlier scans (scrubbing), physiological noise correction (CompCor), and temporal filtering – condition-dependent connectivity effects were inferred from subject-level generalized psychophysiological interaction (gPPI) contrast estimates rather than from raw time-series correlations. This GLM-based framework reduces the likelihood that observed PPI effects reflect differences in temporal autocorrelation or spectral properties across regions rather than genuine task-dependent interactions.

For ROI-to-ROI analyses, the pulvinar divisions, MD, and LGN were defined as seed regions, and connectivity was assessed with target regions including the dACC, sgACC, vmPFC, amygdala, and hippocampus. At the first level, functional connectivity between ROIs was computed as Pearson correlation coefficients of the denoised BOLD time series, which were then Fisher z-transformed to normalize the distribution. To estimate task-related connectivity, the first eight trials of each condition were analyzed as a block.

At the second level, these Fisher z-transformed connectivity values were entered into group-level GLMs to evaluate condition-dependent differences. Paired t-tests were used to assess differences between conditions, with false discovery rate (FDR, *p < 0.05*) correction for multiple comparisons. To provide robust estimates of mean differences, 10,000 bootstrap resamples were used to calculate the BCa CI.

### Masks

We defined the pulvinar divisions, MD, and LGN nuclei using predefined masks based on the neuroanatomical guidelines of the Automated Anatomical Labelling Atlas(Rolls et al., 2020). We also applied the same atlas guidelines to determine the masks for the two other anatomical regions; the amygdala and hippocampus. The masks for the functional regions were created using Neurosynth (Yarkoni et al., 2011) and the keyword ‘conditioning.’ For each region, we created 8mm spheres around the following identified peak coordinates: vmPFC (MNIxyz = −2, 46, −10), sgACC (MNIxyz = 0, 26, −12), and dACC (MNIxyz = 0, 14, 28).

## Acknowledgments

This work was supported by the National Institute of Mental Health grants R01MH123736, R01MH125198, R33MH111907, R01MH097880, and R01MH097964 to M.R.M.

## Data and Code Availability

The data supporting the findings of this study are part of a larger, actively used database and cannot be publicly released at this time due to ongoing projects. The data underlying the analyses reported in this manuscript are available from the corresponding author upon reasonable request and subject to standard data-use agreements.

All scripts used for preprocessing and analysis, including workflows implemented in SPM12, and CONN, are available upon request to facilitate replication and reuse.

