## Supplementary Figure 2 for "Neural Representation of Associative Threat Learning in Pulvinar Divisions, Lateral Geniculate Nucleus, and Mediodorsal Thalamus in Humans"

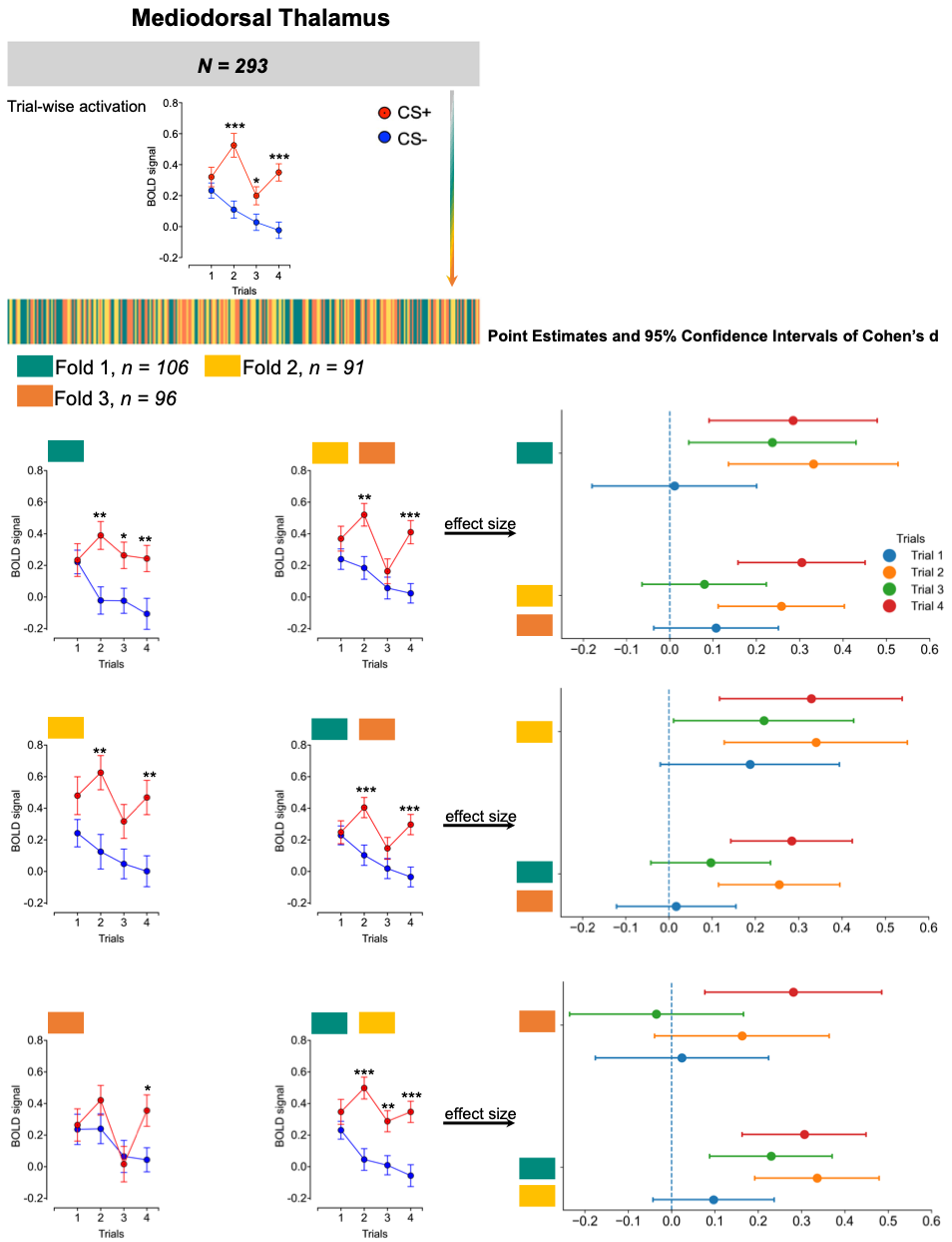
**Supplementary Figure 2**

**Supplementary Figure 2. Activation in the mediodorsal thalamus across cross-validation folds during threat learning.**

Across-validation analysis. The full sample (*N = 293*) was randomly divided into three subsamples (*n_1_ = 106, n_2_ = 91, n_3_ = 96*). For each iteration, we conducted a repeated-measures ANOVA within one subsample and examined the stability of the CS+ vs. CS− difference in the remaining two subsamples combined.
**Top panel:** Mean activation (± SEM) of the overall sample.

**Left panel:** Mean activation (± SEM) across folds.
**Right panel:** Effect size estimates (Cohen’s d) with 95% confidence intervals of the differences between CS+ vs. CS-.
**Note.** False discovery rate (FDR) correction was applied to within-fold comparisons of CS types. *p<0.05, **p<0.01, ***p<0.001
