## Supplementary Figure 3 for "Neural Representation of Associative Threat Learning in Pulvinar Divisions, Lateral Geniculate Nucleus, and Mediodorsal Thalamus in Humans"

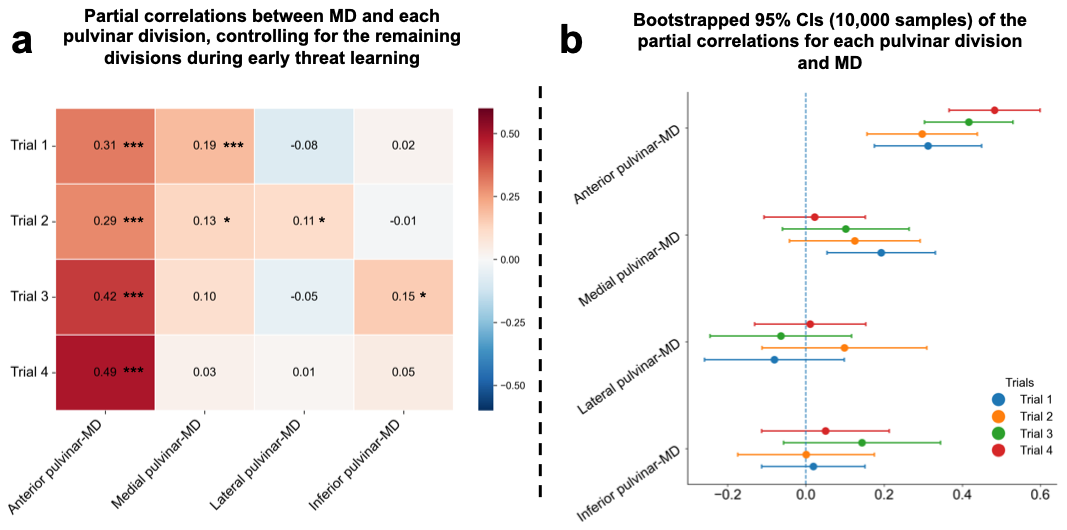

**Supplementary Figure 3.** Partial correlations and bootstrap 95% confidence intervals for associations between the mediodorsal thalamus and pulvinar divisions during threat learning.

(a). Magnitude of the partial correlations between the MD and each pulvinar division, controlling for the remaining pulvinar divisions.

(b) Nonparametric bootstrap 95% confidence intervals (10,000 resamples) of the magnitude of the partial correlations.

MD, mediodorsal thalamus. *p<0.05, **p<0.01, ***p<0.001
