## Supplementary Figure 4 for "Neural Representation of Associative Threat Learning in Pulvinar Divisions, Lateral Geniculate Nucleus, and Mediodorsal Thalamus in Humans"

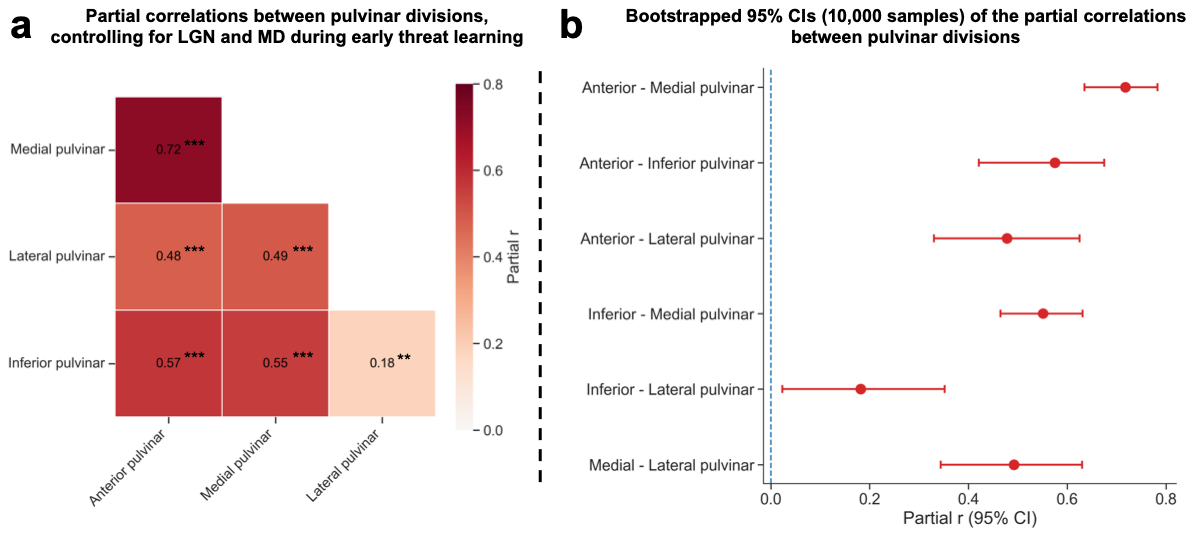

**Supplementary Figure 4.** Partial correlations and bootstrap 95% confidence intervals for associations between pulvinar divisions while controlling for the MD and LGN during threat learning.

(a) Magnitude of the partial correlations between pulvinar divisions, controlling for the MD and LGN.

(b) Nonparametric bootstrap 95% confidence intervals (10,000 resamples) of the magnitude of the partial correlations.

MD, mediodorsal thalamus. LGN, lateral geniculate nucleus. **p<0.01, ***p<0.001
