## Supplementary File 1 for "Neural Representation of Associative Threat Learning in Pulvinar Divisions, Lateral Geniculate Nucleus, and Mediodorsal Thalamus in Humans"

**Results of Repeated-Measures ANOVA for CS+ vs. CS− Differences in the Anterior Pulvinar and Mediodorsal Thalamus**

Two regions—the anterior pulvinar and the mediodorsal thalamus (MD)—showed activation patterns consistent with learning. To evaluate the stability of these effects, we conducted a cross-validation analysis. The full sample (*N = 293*) was randomly divided into three subsamples (*n_1_ = 106, n_2_ = 91, n_3_ = 96*). In each iteration, a repeated-measures ANOVA was conducted within one subsample, and the stability of the CS+ vs. CS− difference was examined in the remaining two subsamples combined. This procedure was repeated such that each subsample served once as the discovery sample and twice as part of the validation sample. Visualization of the means and standard errors of the FDR-corrected CS+ vs. CS− differences is presented in **Supplementary Figures 1–2**. Below, we report the F statistics, n, p values, and effect sizes for the main effects of CS condition.

**Table 1.** Repeated-Measures ANOVA results for the CS effect in the **anterior pulvinar** across cross-validation folds.

|  | **F** | **n** | **p values** | **Effect size (η²_p_)** |
| --- | --- | --- | --- | --- |
| **Overall sample** | 19.47 | 293 | 1.434x10^-5^ | 0.063 |
| **Fold 1** | 10.67 | 106 | 1.46x10^-3^ | 0.092 |
| **Fold 2** | 8.31 | 91 | 4.92x10^-3^ | 0.085 |
| **Fold 3** | 2.22 | 96 | 0.13 | 0.023 |
| **Folds 1+2** | 18.97 | 197 | 2.13x10^-5^ | 0.088 |
| **Folds 1+3** | 11.26 | 202 | 9.43x10^-4^ | 0.053 |
| **Folds 2+3** | 9.59 | 187 | 2.24x10^-3^ | 0.049 |

|  | **F** | **n** | **p values** | **Effect size (η²_p_)** |
| --- | --- | --- | --- | --- |
| **Overall sample** | 41.08 | 293 | 5.83x10^-10^ | 0.123 |
| **Fold 1** | 17.12 | 106 | 7.07x10^-5^ | 0.140 |
| **Fold 2** | 25.14 | 91 | 2.65x10^-6^ | 0.218 |
| **Fold 3** | 3.49 | 96 | 0.06 | 0.036 |
| **Folds 1+2** | 41.791 | 197 | 7.98x10^-10^ | 0.176 |
| **Folds 1+3** | 18.63 | 202 | 2.48x10^-5^ | 0.085 |
| **Folds 2+3** | 23.99 | 187 | 2.09x10^-6^ | 0.114 |

**Table 2.** Repeated-Measures ANOVA results for the CS effect in the **mediodorsal thalamus** across cross-validation folds.
